# Pathogen Clearance-based Correlates of Immunoprotection Reveal Insightful Features of Vaccine Immunity

**DOI:** 10.1101/2022.08.23.504471

**Authors:** Xianbin Tian, Juanjuan Wang, Haoze Chen, Ming Ding, Qian Jin, Jing-Ren Zhang

## Abstract

Vaccination has significantly reduced the incidence of invasive infections caused by several bacterial pathogens, including *Streptococcus pneumoniae, Haemophilus influenzae* and *Neisseria meningitidis*. However, no vaccines are available for many other invasive pathogens. A major hurdle in vaccine development is the lack of functional markers to quantify vaccine immunity in eliminating pathogens in the process of infection. Based on our recent discovery of the liver as the major organ of vaccine-induced clearance of blood-borne virulent bacteria, we here describe a new vaccine evaluation system that quantitatively characterize important properties of effective vaccines in shuffling virulent bacteria from the blood circulation to the liver resident macrophage Kupffer cells (KCs) and sinusoidal endothelial cells (LSECs) in animal models. This system consists of three related correlates or assays: pathogen clearance from the bloodstream, pathogen trapping in the liver, and pathogen capture by KCs/LSECs. The readouts of these correlates were consistently associated with the serotype-specific immunoprotection levels of the 13-valent pneumococcal polysaccharide conjugate vaccine (PCV13) against lethal infections of *S. pneumoniae,* a major invasive pathogen of community-acquired infections in humans. Furthermore, the reliability and sensitivity of these correlates in reflecting vaccine efficacy were verified with whole cell vaccines of *Klebsiella pneumoniae* and *Escherichia coli*, two major pathogens in hospital-acquired invasive infections. This system may be used as cost effective readouts to evaluate the immunoprotective potential of vaccine candidates in the preclinical phase by filling the current technical gap in vaccine evaluation between the conventional *in vitro* approaches (e.g., antibody production and pathogen neutralization/opsonophagocytosis) and survival of immunized animals.

## INTRODUCTION

Sepsis caused by microbial infections of the blood circulation represents an important global health problem and economic burden [1,2]. Approximately 49 million sepsis cases and 11 million sepsis-associated deaths were reported worldwide in 2017, representing 19.7% of all global deaths [2]. Sepsis pathogenesis is initiated by the entry and proliferation of microbial pathogens in the blood circulation, subsequent induction of excessive inflammatory responses, and life-threatening multi-organ dysfunctions [3]. Pathogenic bacteria are the common pathogens of septic infections, which consist of both the Gram-positive bacteria (*Streptococcus pneumoniae*, *Staphylococcus aureus*, *Staphylococcus epidermidis*, and *Enterococcus faecalis*) and Gram-negative bacteria (*Acinetobacter baumannii*, *Escherichia coli*, *Klebsiella pneumoniae*, and *Pseudomonas aeruginosa*) [4].

*S. pneumoniae* (pneumococcus), an encapsulated Gram-positive bacterium residing in the nasopharynx of humans, is an opportunistic pathogen of community-acquired pneumonia, bacteremia, and meningitis, particularly in children under 5 years old and the elderly [5]. There are forms of capsular polysaccharide (CPS)-based vaccines globally licensed to prevent invasive pneumococcal diseases (IPDs) (e.g. bacteremia and meningitis): pneumococcal polysaccharide vaccine (PPV) and pneumococcal conjugate vaccine (PCV) [6]. PPV is the first generation of pneumococcal vaccine that contains T cell-independent CPS antigens with poor immunogenicity in children under two years of ages [7]. PCV represents the next generation of CPS-based vaccine, in which CPSs are chemically linked to carrier proteins (e.g. diphtheria and tetanus toxoids) to activate both B and T lymphocytes to produce IgG antibodies [8]. The global recommendation of PCV has significantly reduced the pneumococcus-associated death worldwide [9].

Vaccination has greatly contributed to the control of many infectious diseases. While programmed immunization has led to global eradication (e.g., smallpox) or nearly eradication (e.g., polio) of a few viral diseases, a number of bacterial diseases have become preventable by vaccination, such as diphtheria (*Corynebacterium diphtheria*), whooping cough (*Bordetella pertussis*), tetanus (*Clostridium tetani*), meningitis (*Neisseria meningitidis*), bacterial pneumonia and sepsis (*S. pneumoniae*, *Haemophilus influenzae*). Except for *S. pneumoniae*, there are no available vaccines for any of the major blood infection pathogens after many unsuccessful efforts in developing vaccines for each of these pathogens. One of the major hurdles in the process of the current vaccine development is the lack of reliable and sensitive immunoprotection correlates to identify any problems in the early steps of vaccine development. Protection efficacy of vaccines relies on two functional aspects of immunization: 1) the features of vaccine-induced antibody and/or T-cell responses, and 2) the potency of vaccine-elicited immunity of pathogen elimination upon pathogen entry [11]. Beyond protection against infections in animal models (e.g., survival rate), the current vaccine evaluation system mostly depends on *in vitro* assessment of vaccine-induced immune responses, such as antibody titer, antibody-mediated neutralization and opsonophagocytosis of pathogen. As an example, the current pneumococcal vaccines are based on measuring antibody level by enzyme-linked immunosorbent assay (ELISA) and antibody-mediated phagocytosis of neutrophil by opsonophagocytic killing assay (OPKA) [12,13]. These parameters don’t inform how vaccine-elicited immunity enable hosts to recognize and clear pathogens in the process of infection. Therefore, there is a great need for reliable and sensitive *in vivo* techniques for functional evaluation of vaccine immunity in the recognition and elimination of invading pathogens in the process of infection.

The liver represents a major organ to capture and eliminate invading microbes from the blood circulation, which are executed by the liver resident macrophage Kupffer cells (KCs) and liver sinusoidal endothelial cells (LSECs) [14]. KCs and LSECs represent approximately 20% and 50% of the liver non-parenchymal cells (NPCs), respectively [15]. KCs are the most dominant resident macrophages in mammals, representing approximately 90% of total resident macrophages in the body [16]. While KCs are specialized for the clearance of large particles (e.g. bacteria) [17], LSECs preferentially clear small particles (e.g. viruses) [18]. Our recent studies have discovered that the liver is the major immune organ to execute vaccine-activated capture and killing of pathogenic bacteria in the bloodstream of mice [19,20]. In particular, PCV13 activates both KCs and LSECs to rapidly recognize and capture IgG-coated pneumococci in the bloodstream [20].

This study aims to establish an *in vivo* vaccine evaluation system to quantify the functional features of vaccine-activated pathogen clearance from the blood circulation to the liver, based on our recent findings of liver-based vaccine immunity [19,20]. We have developed three related new correlates or assays: pathogen clearance from the bloodstream, pathogen trapping in the liver, and pathogen capture by KCs/LSECs in the liver sinusoids. The reliability and sensitivity of these assays have been tested in other two pathogens and vaccines. Potential applications of this new system in vaccine evaluation and development are discussed.

## RESULTS

### The specific antibody level is incompatible with relative protection efficacy of PCV13

To assess the PCV13-induced immune responses, we first measured CPS-specific IgG antibodies in PCV13-immunized and control mice by enzyme-linked immunosorbent assay (ELISA) because antibody titer has been used as an efficacy marker of pneumococcal polysaccharide conjugate vaccines [22,23]. In particular, we focused on the five high-virulence (HV) serotypes (1, 3, 4, 5, and 6A) covered by PCV13 because the protection of these serotypes with relatively low infection doses highly depends on PCV13 immunization in mice [20], and infection caused by the rest of the eight low-virulence (LV) serotypes (6B, 7F, 9V, 14, 18C, 19F, 19A, and 23F) leads to no death in naïve mice [21]. PCV13 immunization generated antibodies to each of the five serotypes, however, the ELISA results revealed substantial variations in antibody level. The antibody level of the highest serotype 4 is more than twice that of the lowest serotype 6A and the order from high to low is 4, 5, 1, 3, 6A (**Fig. 1A**).

**Figure 1.**
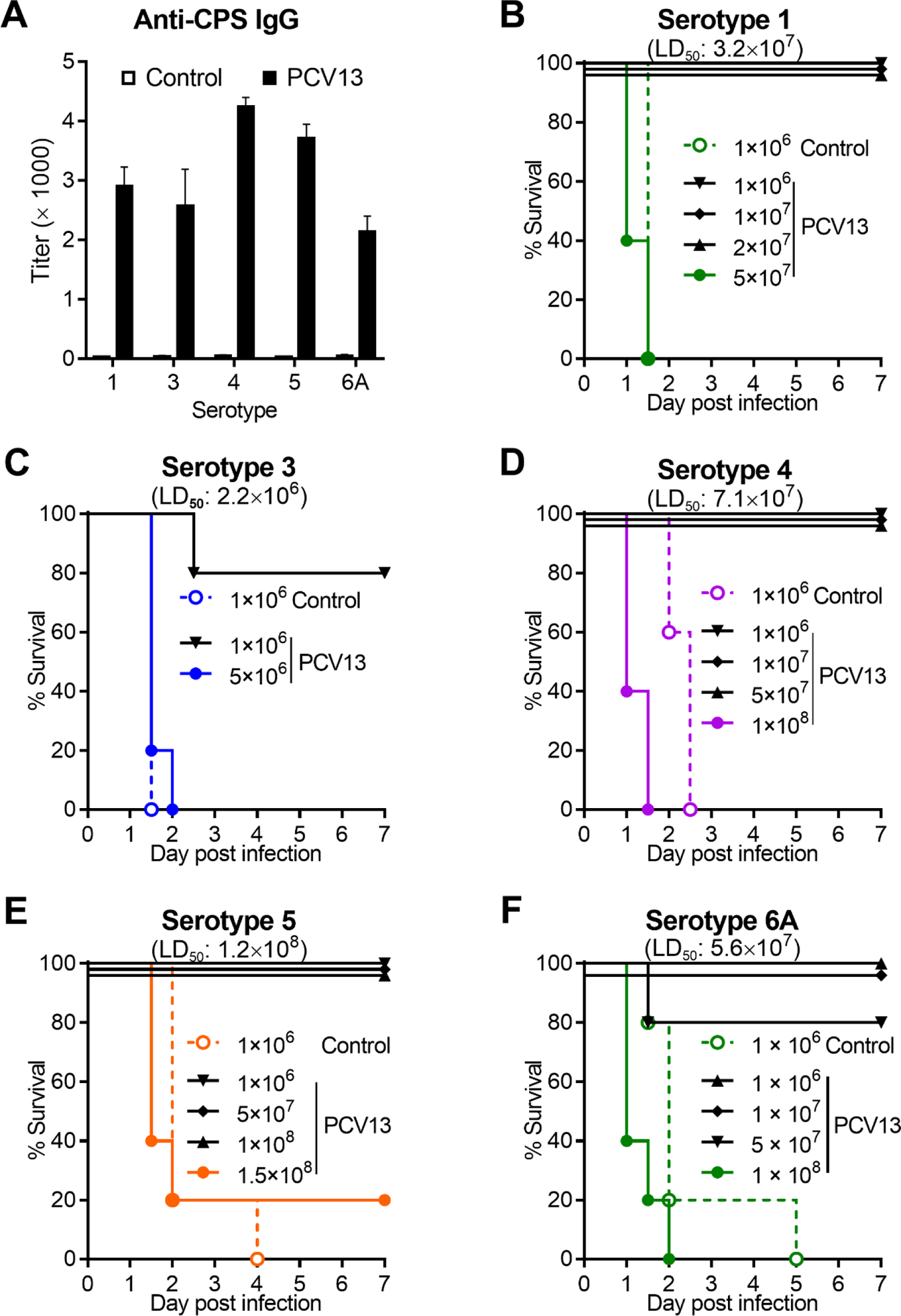
Vaccine-elicited specific antibody level and elevated survival in PCV13-imminzed mice. **(A)**CPS-specific IgG titers of serotype 1, 3, 4, 5, 6A in sera of adjuvant-treated (control) and PCV13-immunized mice as detected by ELISA. n = 3-5. **(B-F)** Survival rate of adjuvant-treated and PCV13-immunized mice i.p. infection with various doses of serotype-1 (**B**), −3 (**C**), -4 (**D**), -5 (**E**) or -6A (**F**) pneumococci. LD_50_ is presented at the top. n = 5. Data presented as the mean values ± standard error of mean (SEM).

To determine if the differences of PCV13-elicited specific antibody of various pneumococcal serotypes reflect protection efficacy against different serotypes of *S. pneumoniae*, we measured the survival of PCV13-immunized mice post intraperitoneal (i.p.) infection. Compared with the full mortality of adjuvant groups post i.p. infection with 10^6^ CFU, PCV13-immunized mice conferred a full protection against serotype 1 (**Fig. 1B**), 4 (**Fig. 1D**), 5 (**Fig. 1E**), and 6A (**Fig. 1F**), and 80% protection against serotype 3 (**Fig. 1C**). We further determined the upper limits of PCV13 immunization against each serotype with higher levels of inoculum. The vaccination conferred the strongest protection against lethal infection of serotype-4, -5, and -6A pneumococci, with a 50% lethal dose (LD_50_) of 7.1 × 10^7^ CFU (**Fig. 1D**), 1.2 × 10^8^ CFU (**Fig. 1E**), and 5.6 × 10^7^ CFU (**Fig. 1F**), respectively. By comparison, the immunized mice were much more susceptible to infection of serotype-3 pneumococci with a LD_50_ of 2.2 × 10^6^ CFU (**Fig. 1C**), and, to a less extent, to that of serotype-1 pneumococci (LD_50_ = 3.2 × 10^7^ CFU) (**Fig. 1B**). Overall comparison revealed a dramatic 54.5-fold difference in LD_50_ between the best (serotype 5) and worst (serotype 3) protected categories. It is noted that the best and worst protected serotypes were not the serotype 4 and 6A with highest and lowest antibody level, respectively. The order of protection efficacy induced by PCV13 immunization against infection of the HV serotypes (5, 4, 6A, 1, and 3) is not the same as the order of antibody level (4, 5, 1, 3, and 6A). The uneven protection efficacy of various serotypes in PCV13-immunized mice strongly suggests that the different antigens in PCV13 elicit variable levels of functional immunity which could not be explained by specific antibody level. These results show that the antibody level is incompatible with relative protection efficacy of PCV13.

### The clearance rate of blood-borne pneumococci predicts the protection efficacy of PCV13

Our recent studies have shown that effective clearance of bacteria from the bloodstream in the early phase of septic infections is essential for host survival [21], and efficient vaccination enhances the pathogen clearance of certain encapsulated high-virulence bacteria that otherwise persist in the blood circulation [19,20]. To assess the direct relationship between the vaccine-elicited pathogen clearance rate and host survival, we compared the PCV13-induced bacterial clearance of different vaccine-covered serotypes of pneumococci from the bloodstream. We monitored the dynamic of bacteremia in immunized and control mice during the first 30 min post intravenous (i.v.) infection of pneumococci. In the adjuvant group, all of the five representative strains retained abundantly in blood circulation within 30 min post i.v. infection of 10^6^ CFU (**Fig. 2A**). In contrast, the circulating pneumococci were cleared quickly from the bloodstream even to undetectable level in PCV13-immunized mice at 30 min post infection (**Fig. 2B**). Consistent with our previous findings of serotype-3 and -4 pneumococci [20], this result demonstrates that PCV13 induces fast pneumococcal clearance of all the five HV serotypes from the bloodstream.

**Figure 2.**
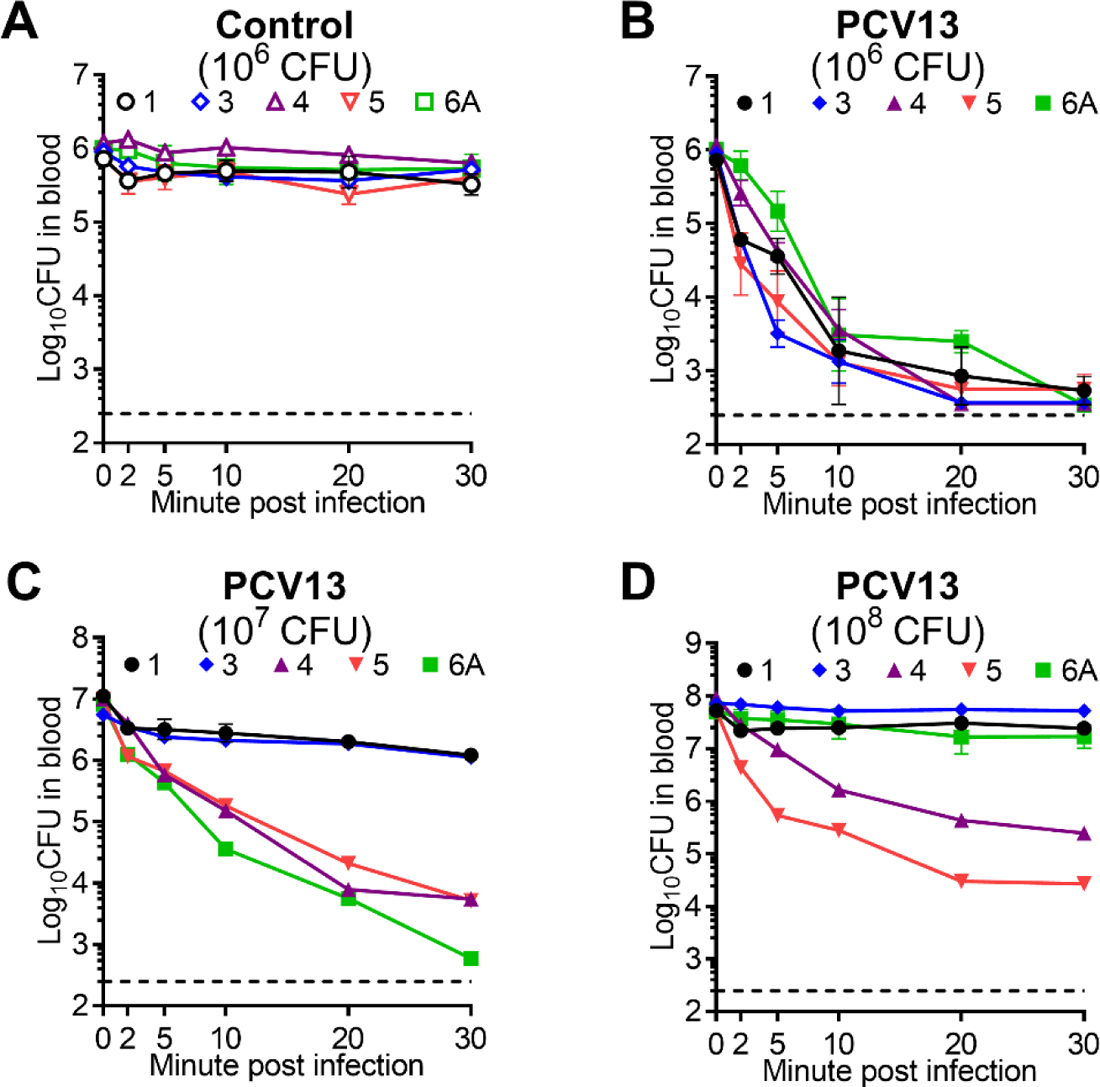
Vaccine-activated pathogen clearance from the circulation in PCV13-imminzed mice. **(A)** Bacteremia levels of adjuvant-treated mice i.v. infected with 10^6^ CFU of serotype-1, -3, -4, -5, -6A pneumococci in the first 30 min. n = 3. **(B-D)** Bacteremia level of PCV13-immunized mice i.v. infected with 10^6^ (**B**), 10^7^ (**C**) or 10^8^ (**D**) CFU of serotype-1, -3, -4, -5, -6A pneumococci in the first 30 min. n = 3. Dotted line indicates the limit of detection. Data presented as the mean values ± standard error of mean (SEM).

Previous studies have indicated that pneumococcal conjugate vaccines PCV7/PCV13 confer relatively poor protection against invasive disease caused by serotype 3 in humans [24–26]. We also found that PCV13 immunization rendered mice uneven ability to combat pneumococcal infection caused by different serotypes (**Fig. 1**). We next tested if pathogen clearance rate could identify this functional difference using higher challenge doses. In line with the different impacts of PCV13 on survival of the five serotypes, PCV13-immunized mice rapidly cleared 10^7^ CFU of serotype-4, -5, and -6A pneumococci from the circulation with clearance half time (CT_50_) of 1.5, 0.7, and 0.8 min, respectively. In sharp contrast, serotype-1 and -3 pneumococci were still largely circulating in the blood of the immunized mice with marginal reduction in bacteremia level post i.v. inoculation with 10^7^ CFU (**Fig. 2C**). With a much higher infection dose (10^8^ CFU), the immunized mice still rapidly cleared 99% inoculum of serotype-5 and -4 pneumococci with CT_50_ of 0.7 and 1.5 min, respectively (**Fig. 2D**), but were much less capable of effectively clearing the other 3 serotypes at this infection dose, with 58%, 74%, and 32% of serotype-1, -3, and -6A inoculum being still detected in the circulation at 30 min, respectively. Overall, these experiments have ranked the functional capacity of PCV13 immunization in clearing the five HV serotypes in the following order: 5, 4, 6A, 1, and 3. The order of blood pathogen clearance rate is coincidence of that of protection efficacy induced by PCV13 immunization. Taken together, these results demonstrate that the PCV13-elicited pathogen clearance rate in the bloodstream is highly predictive of the protection efficacy.

### Hepatic trapping of blood-borne pneumococci correlates the protection efficacy of PCV13

Our recent studies have found that the liver is the major executive organ for vaccine-induced trapping of high-virulence encapsulated *S. pneumoniae* [20] and *Klebsiella pneumoniae* serotypes [19] from the blood circulation. To determine the potential of vaccine-activated hepatic pathogen trapping as a functional correlate for vaccine protection efficacy, we detected the bacterial CFU count in five major organs and blood as previously described [20]. Consistent with the poor clearance of the five HV serotypes in mice of adjuvant group, these animals showed marginal pathogen trapping in the liver for serotypes 1, 3, 4, 5, and 6A post i.v. infection with 10^6^ CFU (**Fig. 3A**). In contrast, virtually all of the viable pneumococci in PCV13-immunized mice were detected in the liver, with near sterilization of the blood circulation, at 30 min post i.v. infection with 10^6^ CFU (**Fig. 3B**). Comparison of the total CFU counts between PCV13 and adjuvant groups revealed significantly lower levels of bacterial burden in vaccinated mice, which reflected vaccine-activated pathogen killing in the liver [20], and the full protection of vaccinated mice against infection with 10^6^ CFU of the five HV serotypes (**Fig. 1**). However, PCV13-immunized mice displayed a serotype-specific pattern of vaccine-activated pathogen trapping in the liver at higher levels of infection dose. Although the majority of viable serotype-1 (77%), -3 (72%), -4 (84%), -5 (91%) and -6A (86%) pneumococci were still found in the livers of PCV13-immunized mice at 30 min post i.v. infection with 10^7^ CFU, serotype-1 (18%) and -3 (20%) pneumococci escaped hepatic capture and remained in the bloodstream (**Fig. 3C**). The vaccine-induced hepatic trapping of the five serotypes is ranked from the highest to the lowest as: 5, 6A, 4, 1 and 3. Relatively poorer hepatic trapping of serotypes 1 and 3 in the immunized mice was also reflected by relatively higher levels of total bacteria detected for these two serotypes at 30 min. When infection dose was increased to 10^8^ CFU, relative proportion of hepatic pathogen trapping was in the order as: 5 (97%), 4 (96%), 6A (21%), 3 (19%), and 1 (3%) and serotype-specific variation in vaccine-induced hepatic pathogen trapping became more dramatic (**Fig. 3D**). These results show that vaccine-activated hepatic trapping of blood-borne pneumococci strongly correlates with the protection efficacy of PCV13.

**Figure 3.**
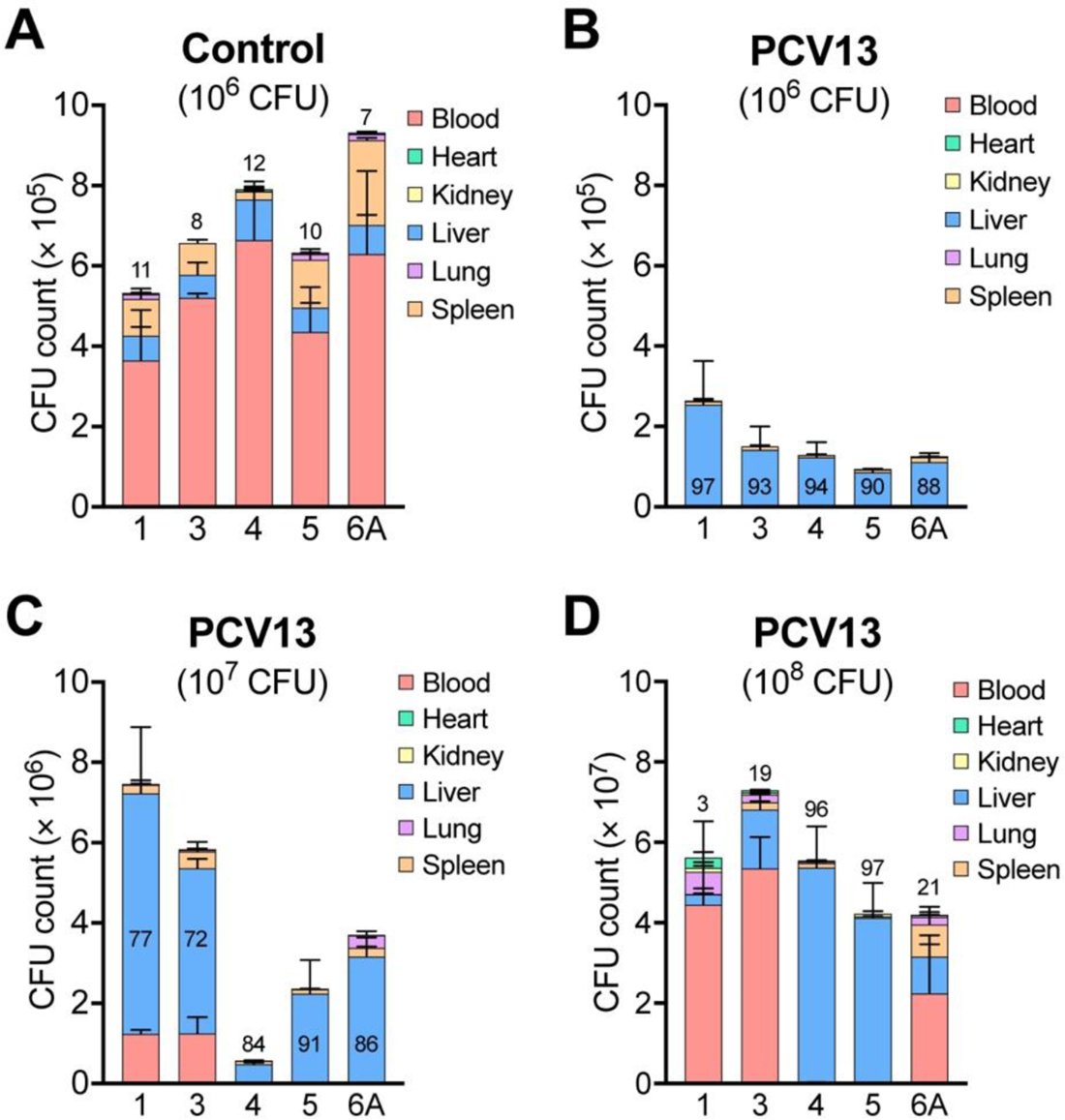
Vaccine-activated hepatic trapping of blood-borne pneumococci in PCV13-immunized mice. **(A)** Proportional bacterial burden in the blood and each of the five major organs of adjuvant-treated mice at 30 min post i.v. infection with 10^6^ CFU of serotype-1, -3, -4, -5 and -6A pneumococci. Relative proportion of hepatic bacteria is presented in or at the top of each bar as a percentage of total bacteria detected in each mouse. n = 3. (**B-D**) Proportional bacterial burden in the blood and five major organs of PCV13-immunized mice at 30 min post i.v. infection with 10^6^ (**B**), 10^7^ (**C**) or 10^8^ (**D**) CFU of serotype-1, -3, -4, -5 and -6A pneumococci. Relative proportion of hepatic bacteria is presented in each bar as in (A). Data presented as the mean values ± standard error of mean (SEM).

### Pathogen capture by liver immune cells indicates the protection efficacy of PCV13

The vaccine-activated dual-track bacterial capture by liver resident macrophages Kupffer cells (KCs) and sinusoidal endothelial cells (LSECs) has been found as the cellular basis of vaccine-elicited pathogen trapping in the liver [20,21]. To assess the potential of cellular capture of circulating pathogens as a functional correlate for vaccine protection efficacy, time-lapse intravital microscopy (IVM) imaging was used to visualize and quantify cellular capture of serotype-1, -3, -4, -5 and -6A pneumococci post i.v. infection with 5 × 10^7^ CFU in the liver sinusoids of PCV13-immunized and control mice. While IVM showed that the HV pneumococci mostly passed the vasculature of control mice (**Fig. 4A, Movies 1-5, left panel**), but were rapidly tethered mainly onto LSECs and less on KCs in immunized mice (**Fig. 4A, Movies 1-5, right panel**). In agreement with the serotype-specific clearance rate induced by PCV13 (**Fig. 2D**), quantification of immobilized bacteria per field of view (FOV) revealed the highest numbers of total captured bacteria for serotypes 4 (110) and 5 (87) as compared with fewer bacteria for serotypes 1 (64), 3 (61) and 6A (74) (**Fig. 4B**). The real-time imaging of vaccine-activated pathogen capture by the liver immune cells in live mice has provided immune cell-based data that are consistent with the protection efficacy of PCV13.

**Figure 4.**
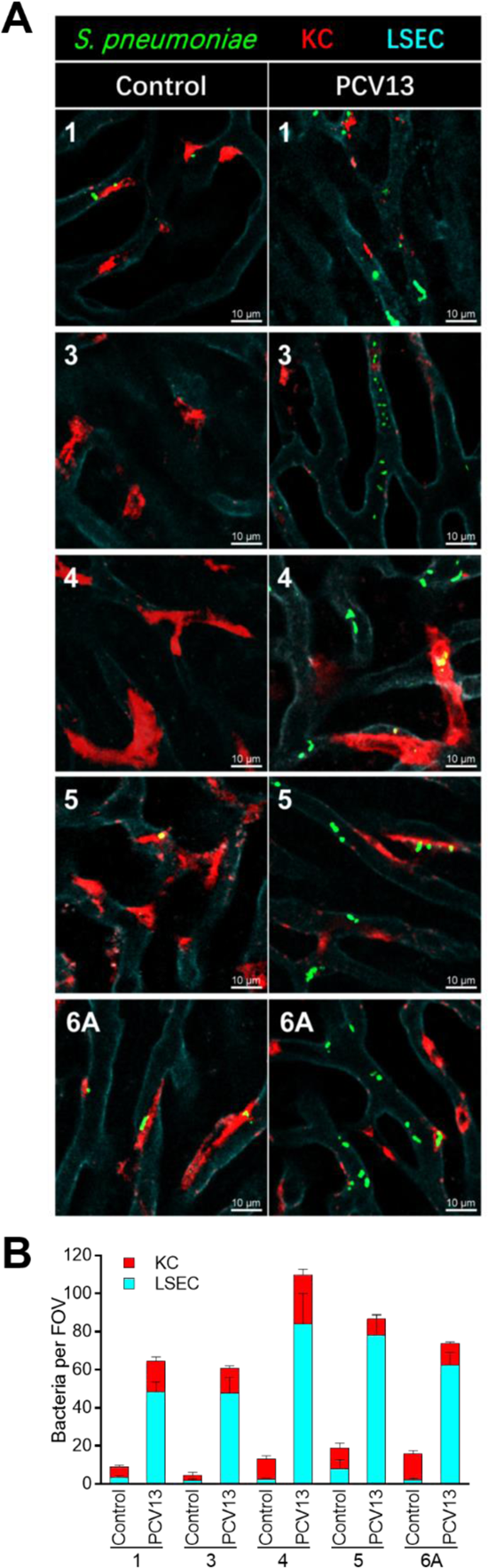
Vaccine-activated pathogen capture by LSECs and KCs in PCV13-immunized mice. **(A)** Representative IVM images of pneumococci (green), Kupffer cells (KCs, red) and liver sinusoid endothelial cells (LSECs, cyan) in the liver sinusoids of adjuvant-treated (control, left panel) or PCV13-immunized (right panel) mice 10 min after i.v. infection with 5 × 10^7^ CFU of serotype-1, -3, -4, -5 and -6A pneumococci. n = 2. The process of bacterial capture is demonstrated in Movies 1-5. **(B)** Quantitation of bacteria immobilized on LSECs and KCs of adjuvant-treated (control) or PCV13-immunized mice in at least five random fields of view (FOV). Data presented as the mean values ± standard error of mean (SEM).

### The *in vivo* immunoprotection correlates reveal limited efficacy of whole cell vaccines

We finally determined whether our PCV13-based *in vivo* vaccine evaluation system could characterize the functional features of other vaccines beyond *S. pneumoniae.* This potential was first tested with an inactivated whole cell vaccine (WCV) for a hypervirulent *Klebsiella pneumoniae* (capsular serotype K2), a Gram-negative bacterium and a major pathogen of blood infections in humans [27]. Our initial trial showed that the WCV confers a full protection against lethal infection of the same *K. pneumoniae* strain (**Fig. 5A**). While all of naïve mice succumbed to infection in the first 36 hr post i.p inoculation with 10^6^ CFU of bacteria, the group of immunized mice completely survived the same challenge. Further analysis revealed that the vaccinated mice gained the ability to clear *K. pneumoniae* from the blood circulation in the first 30 min post i.v. inoculation, with a CT_50_ of 0.6 min, whereas the pathogen were mostly circulating in the bloodstream of naïve mice (**Fig. 5B**). Quantitation of bacteria in the liver and other major organs uncovered 92% of detectable *K. pneumoniae* in the livers of vaccinated mice as compared with 28% hepatic bacteria in naïve mice at 30 min post i.v. infection (**Fig. 5C**). The vaccine-activated clearance of blood-borne *K. pneumoniae* was also manifested by virtual sterilization of the bloodstream in vaccinated mice at 30 min as compared with severe bacteremia in naïve mice (43% of total bacteria in blood). IVM imaging revealed significantly enhanced capture of circulating bacteria by KCs but not LSECs in the sinusoids of vaccinated mice (**Fig. 5D** and **5I**), which is consistent with relatively low protection capacity of the WCV (full mortality at higher infection dose of 10^7^ CFU, **Fig. 5A**). We further confirmed the reliability of the *in vivo* immunoprotection correlates with inactivated WCV of a virulent *Escherichia coli* K1 strain, a capsular serotype commonly causing human blood infection and neonate meningitis[27]. The i.p. infection with a lethal dose of the same *E. coli* strain led to 100% mortality in naïve mice but a complete protection in the vaccinated mice (**Fig. 5E**). In line with the immunoprotection capacity, WCV immunized mice also showed rapid clearance of the pathogen from the circulation (**Fig. 5F**), effective pathogen capture in the liver (**Fig. 5G**) by KCs (**Fig. 5H** and **5J**), as evidenced by dominant proportion of the detectable bacteria in the liver (81%) and virtual blood sterilization of each immunized mouse. It should be mentioned that the KCs of *E. coli* WCV-immunized mice were much more capable of pathogen capture (94 bacteria per FOV) than the counterparts of *K. pneumoniae-*WCV (13 bacteria per FOV), suggesting that the former is more effective. This notion is supported by more significant pathogen killing in the *E. coli* WCV-immunized mice, as indicated by more significant reduction in the total bacteria in *E. coli* WCV-immunized mice (53% of the total bacteria) than the *K. pneumoniae* counterparts (104% of the total bacteria). Lastly, IVM imaging did not revealed any significant enhancement of pathogen capture by LSECs by either *K. pneumoniae* WCV (**Fig. 5I**) or *E. coli* WCV (**Fig. 5J**), which is in sharp contrast to the dominant role of LSECs in PCV13-immunized mice (**Fig. 4**). Based on the extreme importance of LSECs in upgrading vaccine protection potency [20], the failure of WCV in activating pathogen capture of LSECs indicated the limited efficacy of the whole cell vaccines as compared with the licensed PCV13. In conclusion, these data have confirmed the reliability of our *in vivo* immunoprotection correlates in quantifying the functional features and potency of vaccine-activated immunity.

**Figure 5.**
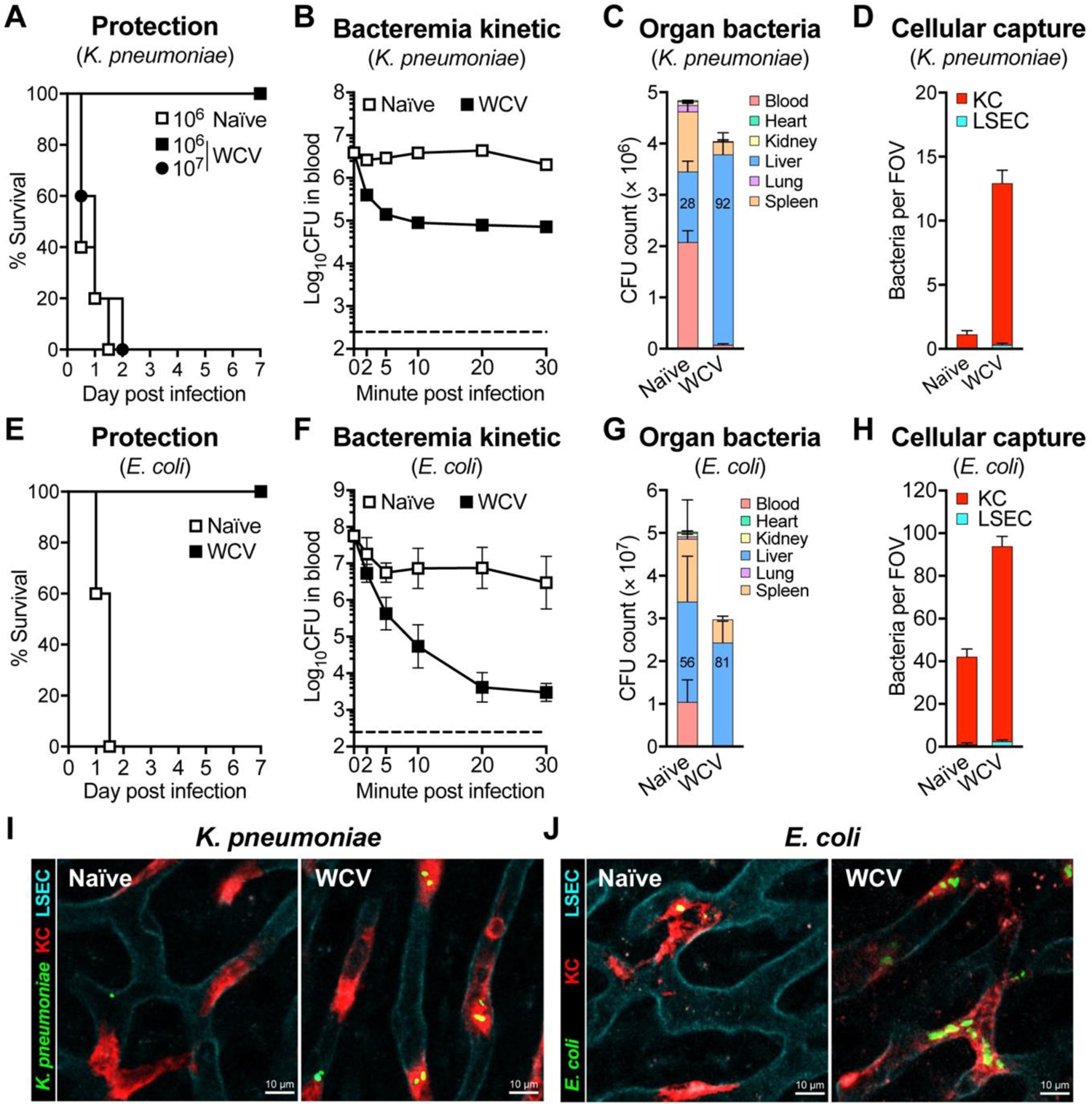
Vaccine-activated pathogen clearance, trapping in the liver and capture by Kupffer cells in whole cell vaccine (WCV)-immunized mice. **(A-D)** Functional features of *K. pneumoniae* WCV. Host survival (**A**), bacteremia in the first 30 min (**B**), bacterial burden in blood and major organs at 30 min (**C**), and bacterial capture by LSECs and KCs (**D**) post infection of *K. pneumoniae* strain TH13044 (serotype K2) in naïve or WCV*-*immunized mice. Infections were performed with the following doses and routes: 10^6^ or 10^7^ CFU (i.p.) for (A), 5 × 10^6^ CFU (i.v.) for (B-C), 5 × 10^7^ CFU (i.v.) for (D). n = 5 for (A), 3 for (B-C), and 2 for (D). (E-H) Functional features of *K. pneumoniae* WCV. Host survival (**E**), bacteremia (**F**), bacterial burden in blood and major organs (**G**), and bacterial capture (**H**) post infection of *E. coli* strain TH14512 (serotype K1) in naïve or WCV*-*immunized mice. Infections were performed with the following doses and routes: 5 × 10^7^ CFU (i.p.) for (E), 10^8^ CFU (i.v.) for (F-G), 5 × 10^7^ CFU (i.v.) for (H). n = 5 for (E), 3 for (F-G), and 2 for (H). **(I-J)** Representative IVM images of the liver sinusoids in naive (left panel) or WCV-immunized (right panel) mice 10 min after i.v. infection with 5 × 10^7^ CFU of *K. pneumoniae* **(I)** and *E. coli* **(J)** as in Fig. 4A. The process of bacterial capture is demonstrated in Movies 6-7. Data presented as the mean values ± standard error of mean (SEM).

## DISCCUSSION

Immunoprotection efficacy is the core function of all vaccines, regardless the natures of pathogenesis and immunization. However, the current vaccine evaluation system almost entirely depends on *in vitro* features of vaccine-induced responses beyond impact on host disease severity and/or survival, such as antibody titer and antibody-mediated neutralization and/or opsonophagocytosis [22,28,29]. The dynamics of vaccine-induced *in vivo* actions toward pathogen elimination in the process of infection have remained as a “black hole”. In this study, we have used three different vaccines (one PCV13 and two WCVs) for three pathogens (*S. pneumoniae*, *K. pneumoniae* and *E. coli*) to characterize an *in vivo* vaccine evaluation system, based on our recent discoveries of the liver as the major immune organ to capture and kill highly virulent bacteria from the blood circulation in vaccinated mice [19,20]. This system includes three related vaccine-activated functions in the defense against invasive infection: pathogen clearance from the bloodstream, trapping in the liver, and cellular capture by KCs and/or LSECs. These correlates have thus bridged our knowledge gap between vaccine-induced antibody response and immunoprotection. Despite that many effective vaccines have been developed by conventional approaches, many important infectious diseases remain unpreventable by vaccination. These functional features of vaccine immunity will not only upgrade our understanding of vaccine biology, but may also empower future vaccine development, particularly for those invasive pathogens that are able to survive in the blood circulation by circumventing innate capture by Kupffer cells (encapsulated bacteria, fungi, and protozoa) and LSECs (viruses).

We have clearly shown that the kinetics of pathogen decay in the bloodstream of vaccinated mice accurately reflect the immunoprotection level of vaccines (survival rate). The pneumococcal serotypes with faster clearance rates or lower CT_50_ values at high infection doses (serotypes 4, 5 and 6A) in PCV13-immunized mice consistently exhibited stronger immunity to lethal challenge or higher LD_50_ values (**Figs. 1 and 2**). Conversely, the serotypes with higher CT_50_ counts also displayed lower LD_50_ values (serotypes 1 and 3). Furthermore, *E. coli* WCV-immunized mice that displayed faster pathogen clearance in the bloodstream than the *K. pneumoniae* counterparts also showed more potent immunoprotection (**Fig. 5**). Vaccine-activated removal of invading bacteria from bloodstream has been commonly used as a parameter for vaccine evaluation [20,30,31]. However, to the best of our knowledge, this is the first report of demonstrating quantitative relationship between vaccine-activated blood pathogen clearance and immunoprotection. An advantage of this parameter is its suitability in repetitive sampling of the blood by retroorbital bleeding without sacrificing the animals. Along with the dynamic detection of blood bacteria at multiple time points post i.v. infection, typically within 30 min, both the clearance proportion and rate of circulating bacteria can be measured (**Fig. 2**), which makes it more sensitive and quantitative.

Our data have shown that the vaccine-activated pathogen trapping in the liver is well suited as a functional correlate for immunoprotection potency of vaccines against blood-borne bacteria. Serotypes 4, 5 and 6A with the superior immunoprotection or high LD_50_ values in PCV13-immunzed mice displayed higher percentages of total bacteria in the liver than serotypes 1 and 3, which exhibited the inferior immunoprotection and hepatic pathogen trapping (**Fig. 3**). Moreover, we have used two WCVs to confirm that vaccine-activated hepatic pathogen trapping can be used to evaluate the immunoprotection potency of vaccines beyond the polysaccharide conjugate vaccines. *E. coli* WCV was superior to the *K. pneumoniae* WCV in both hepatic pathogen trapping and immunoprotection efficacy (**Fig. 5**). The liver has been recognized as a major organ to clear blood-borne microbes for over six decades [32–37]. Our recent studies have uncovered the essential role of the liver as the organ for the execution of vaccine-activated immunity against blood infections of virulent bacteria [19,20], which has changed our conventional view that vaccine-activated immunity eliminates blood-borne bacteria by circulating phagocytes [6,38].

Consistent with the abovementioned causal relationship between hepatic pathogen trapping and immunoprotection efficacy, this study has demonstrated that cellular pathogen capture by KCs and LSECs in the liver sinusoids is a reliable and sensitive indicator of vaccine immunoprotection. This relies on the state-of-art non-invasive intravital imaging technology to visualize the process of pathogen capture by KCs and LSECs in the liver sinusoidal vasculatures. This technique can provide multiple facets of functional information on vaccine immunity: overall pathogen capture and ratio of KC- and LSEC-captured bacteria. As manifested by significant elevation of PCV13 in FOV value (**Fig. 4**), the vaccine activating stronger pathogen capture by both KCs and ECs confers more potent immunoprotection. However, as indicated in our recent work [20], LSEC-mediated pathogen capture is an accurate correlate of the elite vaccine immunity. LSEC-dominant pathogen capture in the liver accompanies the potent immunoprotection in PCV13-immunized mice. In sharp contrast, the mice immunized with two WCVs showed only pathogen capture by KCs and relatively weaker immunoprotection efficacies. Along this line, effective activation of LSEC’s pathogen capture will be the direction for the development of potent vaccines for *E. coli*, *K. pneumoniae* and other common pathogens of blood infections in humans.

This study has uncovered important insights into the spatial and technical venues for future improvement of polysaccharide conjugate vaccines. While previous epidemiological surveys have reported relatively poor protection of pneumococcal conjugate vaccines against serotype-3 pneumococci in humans [24-26,39], the mechanisms behind the functional differences between serotype 3 and other types have remained undefined. Our data in mouse blood infection model have shown that, despite comparable levels of PCV13-induced serotype-specific IgG antibodies for the five serotypes, PCV13-induced immunoprotection against serotype-3 is strikingly inferior to other serotypes (e.g., LD_50_ values). Accordingly, our *in vivo* vaccine evaluation system reveals that PCV13 induces much poorer levels of blood pathogen clearance, hepatic pathogen trapping and cellular pathogen capture in the liver. Because vaccine-induced IgG antibodies, particularly IgG1 subtype, define the pathogen binding from KCs to LSECs in the liver [20], our data indicate that PCV13 induces a much lower level of protective antibodies to the serotype-3 capsule, which cannot be identified by measuring the total antibodies.

In summary, our *in vivo* vaccine evaluation system can not only reliably and sensitively distinguish the functional features of vaccines, but also provide functional guidance to future vaccine design and evaluation. The three assays of this new vaccine evaluation system can be directly applicable to the evaluation and development of new vaccines for bacteria, fungi and parasites that cause systemic infections, particularly the encapsulated bacteria that are refractory to phagocytic clearance in the liver (e.g., *E. coli* and *K. pneumoniae*). With certain technical modifications, our findings may also be used for evaluation of vaccine candidates for viral pathogens that depend on viremia for dissemination and vector-borne transmission.

## METHOD AND MATERIAL

### Bacteria strains and cultivation

All the strains of *S. pneumoniae*, *E. coli* and *K. pneumoniae* used in this study are listed in **Table S1**. Pneumococcal strains were cultured in Todd-Hewitt broth (THB) (Oxoid) with 0.5% yeast extract or on tryptic soy agar (TSA) (BD Difco) plates with 5% defibrinated sheep blood at 37℃ with 5% CO_2_ as described [21]. *E. coli* and *K. pneumoniae* were cultured in Luria-Bertani (LB) broth at 37℃ with shaking as described [19].

### Vaccines and immunoprotection

PCV13 used in this study have been approved by the Chinese National Medical Products Administration (NMPA). C57BL/6J mice and CD1 mice were obtained from Beijing HFK Bioscience (Beijing, China) and Vital River (Beijing, China), respectively, and kept in SPF facilities at Laboratory Animal Resources Center of Tsinghua University. All animal experiments were conducted according to animal protocols approved by the Institutional Animal Care and Use Committee at Tsinghua University. Immunization with PCV13 was carried out in 5-week-old C57BL/6J mice as previously described [20]. 50 μl (1/10 dose of human) of PCV13 or adjuvant (aluminum phosphate) was intramuscularly (i.m.) injected. Two boost immunizations were carried out by the same procedure 14 and 28 days later. Immunization with the whole cell vaccines of *E. coli* and *K. pneumoniae* was carried out in 5-week-old CD1 mice as previously described[43, 44]. Briefly, 10^8^ colony forming unit (CFU) of *E. coli* strain TH14512 (K1) or *K. pneumoniae* strain TH13044 (K2) were treated with 0.5% formalin in phosphate-buffered saline (PBS) at 4°C overnight, washed with PBS by centrifugation, resuspended in 475 μl PBS, and mixed with 25 μl of Adju-Phos® adjuvant (InvivoGen) before subcutaneous (s.c.) immunization. The same procedure was repeated once 7 days later. Immune sera were collected by retro-orbital bleeding 2 weeks after final immunization.

Immunoprotection was assessed by intraperitoneal (i.p.) infection with various doses of bacteria in 100 μl PBS 3 weeks post final immunization as described [20]. The animal survival was observed every 12 hrs within 7 days post infection. The 50% lethal dose (LD_50_) was calculated by LD_50_ calculator (Quest GraphTM, AAT Bioquest, Inc).

### Bacterial clearance from the bloodstream

The clearance of *S. pneumoniae*, *E. coli* and *K. pneumoniae* from the bloodstream was evaluated by assessing kinetics of bacteremia retro-orbital bleeding and CFU plating on TSA blood plates at various time points in the first 30 min post intravenous (i.v.) inoculation of bacteria in 100 μl PBS as described previously [20]. Clearance time for half of the inoculum (CT_50_) values were obtained as described [21].

### Bacterial trapping in the liver

Bacteria trapped in the liver and other major organs (heart, kidney, lung and spleen) were determined as described [20]. Briefly, bacterial burdens in the blood, heart, kidney, liver, lung and spleen of each mouse were measured by CFU plating at 30 min post infection and presented as CFU count for each organ. Bacteria trapped in the liver are presented as CFU count or percentage of bacteria detected in the liver out of the total viable bacteria detected in the blood and five major organs.

### Cellular capture of bacteria by liver immune cells

Bacterial capture by KCs and LSECs was assessed by real-time intravital microscopy (IVM) imaging of mouse liver with inverted confocal laser scanning microscopy as described [45]. The LSECs and KCs were labelled by i.v. injection of 2.5 μg AF594 anti-CD31 (BioLegend) and AF647 anti-F4/80 (BioLegend) antibodies, respectively, 30 min before infection. 5 × 10^7^ CFU of FITC-labeled bacteria were i.v. inoculated to track the immobile bacteria captured in the liver sinusoids. FITC labeling was accomplished by incubating 10^8^ CFU of bacteria with 100 μl PBS containing 1 μg FITC (Sigma-Aldrich) for 30 min at room temperature in dark. Images were acquired with Leica TCS SP8 confocal microscope using 10×/0.45 NA and 20×/0.80 NA HC PL APO objectives. The microscope was equipped with Acousto Optics without filters. Fluorescence signals were detected by photomultiplier tubes and hybrid photo detectors (600 × 600 pixels for time-lapse series and 1,024 × 1,024 pixels for photographs). Three laser excitation wavelengths (488, 585, and 635 nm) were employed by white light laser (1.5 mw, Laser kit WLL2, 470–670 nm). Real-time imaging was monitored for at least 1 min post infection. More than five random fields of view (FOV) at 10 min post infection were selected to quantify the bacteria captured by LSECs and KCs.

### Enzyme-linked immunosorbent assay (ELISA)

CPS-specific IgG was quantified by enzyme-linked immunosorbent assay (ELISA) as described [20,46]. Briefly, 96-well plates were coated with capsular polysaccharides and then blocked by 5% non-fat dry milk (BD Difco). The mouse serum was serially diluted with PBS and added into CPS-coated wells for two-hour incubation at 37°C. Bound antibodies were detected by incubation with goat anti-mouse HRP-conjugated IgG (1:2,000, EasyBio, Beijing, China) for 1 h. The optical absorbance at wavelength of 450 nm was measured after 5-minute incubation of TMB substrates (TIANGEN, China) and stopped by the addition of 1 M phosphoric acid.

## Acknowledgments

We thank Tsinghua research platforms for assistance in animal experimentation (Laboratory Animal Research Center) and IVM imaging (Center for Cell Biology).

## Funding

This work was supported by grants from National Natural Science Foundation of China 31820103001, 31530082, 81671972, 31728002 (J.-R.Z.) and 3210010245 (J.W.); China Postdoctoral Science Foundation 2020M680518 (J.W.); Tsinghua-Peking Joint Center for Life Sciences Postdoctoral Foundation (J.W.).

## Author contributions

Conceptualization: X.T., J.W., J.-R.Z.

Experimentation: X.T., J.W., H.C., M.D., Q.J., J.-R.Z.

Methodology: X.T., J.W., H.C., M.D., Q.J.

Investigation: J X.T., J.W., J.-R.Z.

Funding: J.W., J.-R.Z.

Project administration: J.W., J.-R.Z.

Supervision: X.T., J.W., J.-R.Z.

Writing original draft: X.T., J.W., J.-R.Z.

## Competing interests

Patent applications have been filed on the basis of this work.

## Data and materials availability

All data are available in the main text or the supplementary materials.

---

**Table S1.** Bacterial strains used in this study

| Strain ID | Species | Serotype | Source | Region | Reference |
| --- | --- | --- | --- | --- | --- |
| TH2866 | *S. pneumoniae* | 1 | Blood | China | [21] |
| TH865 | *S. pneumoniae* | 3 | Cerebrospinal Fluid | USA | [21] |
| TIGR4 | *S. pneumoniae* | 4 | Blood | Norway | [21] |
| TH2940 | *S. pneumoniae* | 5 | Blood | China | [21] |
| TH870 | *S. pneumoniae* | 6A | Blood | USA | [21] |
| TH13044 | *K. pneumoniae* | K2 | Blood | China | [19] |
| TH14512 | *E. coli* | K1 | Blood | China | [21] |

## SUPLMENATRY MATERIALS

**Movie 1-5. PCV13-elicited pneumococcal capture of high-virulence serotypes in liver sinusoid.** Capture of i.v. inoculated serotype-1 (Movie 1), -3 (Movie 2), -4 (Movie 3), -5 (Movie 4), and -6A (Movie 5) pneumococci (5 × 10^7^ CFU, FITC-labelled, green) from the bloodstream by liver sinusoid endothelial cells (LSECs, cyan) and resident macrophage Kupffer cells (KCs, red) in adjuvant-treated (control) or PCV13-immunized mice. Real-time imaging was visualized for at least 1 minute by intravital microscopy immediately post infection. The playback rate is eight frames per second. The quantitative data were shown in Fig. 4B; selected images in Fig. 4A.

**Movie 6-7. Whole cell vaccine induced capture of *K. pneumoniae* and *E. coli* by KCs**. 5 × 10^7^ CFU of FITC-labelled *K. pneumoniae* strain TH13044 (Movie 6) and *E. coli* strain TH14512 (Movie 7) in naive or WCV-immunized mice were visualized as in Movie 1. The quantitative data were shown in Figs. 5D and 5H; selected images in Figs. 5I and 5J.

## Notes

### Summary of Updates

Original Figure 1 is separated into new Figure 1 and new Figure 2; There are no changes in other figures except original figure 4 and related information are deleted because of lack of significance different; Some spelling mistakes are corrected.

